# Fine-tuning of the PAX-SIX-EYA-DACH network by multiple microRNAs controls embryo myogenesis

**DOI:** 10.1101/2020.02.18.954446

**Authors:** Camille Viaut, Andrea Münsterberg

## Abstract

MicroRNAs (miRNAs), short non-coding RNAs, which act post-transcriptionally to regulate gene expression, are of widespread significance during development and disease, including muscle disease. Advances in sequencing technology and bioinformatics led to the identification of a large number of miRNAs in vertebrates and other species, however, for many of these miRNAs specific roles have not yet been determined. LNA *in situ* hybridisation has revealed expression patterns of somite-enriched miRNAs, here we focus on characterising the functions of miR-128. We show that antagomir-mediated knock-down (KD) of miR-128 in developing chick somites has a negative impact on skeletal myogenesis. Computational analysis identified the transcription factor *EYA4* as a candidate target consistent with the observation that miR-128 and *EYA4* display similar expression profiles. Luciferase assays confirmed that miR-128 interacts with the *EYA4* 3’UTR. Furthermore, *in vivo* experiments suggest that *EYA4* is regulated by miR-128, as *EYA4* expression is derepressed after antagomir-mediated inhibition of miR-128. EYA4 is a member of the PAX-SIX-EYA-DACH (PSED) network of transcription factors. Therefore, we identified additional candidate miRNA binding sites in the 3’UTR of *SIX1/4, EYA1/2/3* and *DACH1*. Using the miRanda algorithm, we found sites for miR-128, as well as for other myogenic miRNAs, miR-1a, miR-206 and miR-133a, some of these were experimentally confirmed as functional miRNA-target sites. Our results reveal that miR-128 is involved in regulating skeletal myogenesis by targeting *EYA4* transcripts and moreover that the PSED network of transcription factors is co-regulated by multiple muscle-enriched microRNAs.

## Introduction

In vertebrates, most of the axial skeleton and all skeletal muscles of the trunk and limbs are derived from somites, transient metameric structures generated along the anterior-posterior axis by segmentation from the pre-somitic mesoderm (PSM) (Christ and Ordahl, 1995). Subsequent specification of cell fates within somites and the differentiation of a somite into sclerotome, dermomyotome and myotome depends on interactions with surrounding tissues, which are the source of extrinsic molecular signals. These signals include WNT proteins derived from the dorsal neural tube and surface ectoderm, bone morphogenetic proteins (BMP) from the lateral plate mesoderm, and Sonic hedgehog (SHH) from the notochord and floor plate of the neural tube (Brent and Tabin, 2002; Christ et al., 2007).

Myogenesis starts in the dermomyotome and requires the commitment of a pool of cells into the skeletal muscle lineage. The first molecular markers characterising myogenic precursors are the paired-box transcription factors PAX3 and PAX7 (Kassar-Duchossoy et al., 2005; Relaix et al., 2005), which support the proliferation and survival of myoblast before differentiation (Buckingham and Relaix, 2007). In both mouse and chicken embryos, PAX3/7 activate and control the expression of the genes encoding myogenic regularoty factors (MRF), such as Myf5 and MyoD1 (Bajard et al., 2006; Maroto et al., 1997; Tajbakhsh et al., 1997; Williams and Ordahl, 1994). Furthermore, mice lacking both *PAX3* and *PAX7* display major defects in myogenesis, suggesting that together these genes are required for normal muscle development (Relaix et al., 2005).

The expression of PAX3/7 is regulated by the activity of members of SIX, EYA and DACH families (Grifone et al., 2005; Heanue et al., 1999). Together these proteins comprise the PSED network, which plays key regulatory roles in the development of numerous organs and tissues such as kidney, ear and muscle (Relaix and Buckingham, 1999). The biochemical interactions and complex feedback loops between PSED members have been dissected (Kumar, 2009). In paraxial mesoderm, the expression of PSED members (PAX1/6/7/9, SIX1/2, EYA2) is upregulated during the transition from presegmented mesoderm to epithelial somites (Mok et al., 2020). In addition, SIX1/4, EYA1/2/4 and DACH1/2 have been shown to initiate myogenesis through activation of the MRF genes, similar to PAX3 and PAX7 (Grifone et al., 2007; Heanue et al., 1999; Maroto et al., 1997; Relaix et al., 2013; Relaix et al., 2005; Spitz et al., 1998; Tajbakhsh et al., 1997; Tapscott, 2005). Thus, the PSED network is upstream of the genetic regulatory cascade that directs dermomyotomal progenitors toward the myogenic lineage.

SIX family transcription factors are characterised by the presence of two conserved domains, a homeodomain (HD) that binds to DNA, and an amino-terminal SIX domain (SD) that interacts with coactivators (EYA) or corepressors (DACH) of transcription. EYA proteins are unique co-transcription factor phosphatases. They comprise a C-terminal EYA domain (ED), responsible for interactions with SIX and DACH, and threonine and tyrosine phosphatase activity, which may inhibit DACH corepressor function (Li et al., 2003; Rayapureddi et al., 2003; Tootle et al., 2003). Furthermore, EYA recruits RNA polymerase II and coactivators, such as CREB-binding protein (CBP), or corepressors, such as histone deacetylase (HDAC), to the SIX complex (Jemc and Rebay, 2007; Li et al., 2003).

It has been shown that microRNAs, small non-coding RNAs that regulate gene expression post-transcriptionally (Bartel, 2004, 2009), are important for embryo myogenesis (Mok et al., 2017). We showed that members of the miR-1/miR-206 and the miR-133 families, which are derived from bi-cistronic primary transcripts, are expressed in the myotome of developing somites where their expression is induced by MRFs (Goljanek-Whysall et al., 2011; Sweetman et al., 2008; Sweetman et al., 2006). Antagomir-mediated knock-down (KD) approaches *in vivo* revealed that miR-206 is crucial for the myogenic progenitor to committed myoblast transition by negatively regulating expression of PAX3. PAX3 is initially expressed throughout the somite (Williams and Ordahl, 1994), subsequently becomes restricted to the dermomyotome and then to the epaxial and hypaxial dermomyotome. PAX3 is finally downregulated as progenitor cells enter myogenesis. Furthermore, we showed that miR-133 and miR-1/206 directly target BAF60a and BAF60b, thereby affecting the subunit composition of the BAF/BRG1 chromatin remodelling complex (Goljanek-Whysall et al., 2014). This is important to stabilize the myogenic differentiation program in developing somites. In addition, miR-133 is involved in regulating Sonic Hedgehog pathway activity via negative regulation of GLI3 repressor and this is required for myogenic fate specification as well as somite epithelialization, proliferation and growth (Mok et al., 2018).

Here we focus on miR-128, which we found enriched in developing somites (Ahmed et al., 2015). miR-128 is intronic and embedded into two distinct genes: *R3HDM1* (R3H domain containing 1) and *ARPP21* (cyclicAMP regulated phosphoprotein 21 kDa) (Bruno et al., 2011), located on chromosome 7 and 2 in chicken. Both miR-128-1 and miR-128-2 precursors generate the identical mature miRNA. First identified in mouse, miR-128 is enriched in brain, during development and in the adult (Lagos-Quintana et al., 2002), similar observations were made in chicken and *zebrafish* (Kapsimali et al., 2007; Xu et al., 2006). Furthermore, during cardiac regeneration in newt, miR-128 regulates the expression of the transcription factor Islet1 (Witman et al., 2013). In chicken, miR-128 expression in the developing heart appears to be limited to a short time-window as it is only seen in stage HH13 embryos (Darnell et al., 2006). As well as being involved in neuronal and cardiac development, miR-128 expression was also detected in adult mouse muscle (Sempere et al., 2004), adult and embryo porcine skeletal muscle (Zhou et al., 2010), and adult and embryo chicken skeletal muscle (Abu-Elmagd et al., 2015; Darnell et al., 2006; Lin et al., 2012). In mouse, the inhibition of insulin receptor substrate 1 (IRS1) by miR-128 leads to inhibition of myoblast proliferation and induction of myotube formation (Motohashi et al., 2013). In addition, miR-128 promotes myotube formation by targeting myostatin (MSTN), a negative regulator of myogenesis and muscle growth and ectopic miR-128 leads to expression of *Pax3/7* and MRFs (Shi et al., 2015). The functions of miR-128 in embryo myogenesis are less well understood. We identify EYA4 as a novel target for miR-128 and show cooperative effects with miR-206. Using luciferase reporter assays we examine the regulation of additional PSED members by miR-128, as well as by other muscle-enriched microRNAs, including miR-1, miR-206 and miR-133. We show that antagomir-mediated knock down (KD) of miR-128 in chick somites inhibits myogenesis and correlates with the deregulation of PSED members, including the derepression of EYA4. Electroporation of EYA4 into chick somites inhibits myogenesis, thus mimicking the miR-128 KD phenotype. Together our findings suggests that microRNA mediated regulation of the PSED network contributes to the fine-tuning of skeletal muscle development in vertebrate embryos.

## Results

### Expression of miR-128 overlaps with Eya4 in the myotome

To analyse the properties of miR-128 during somite development in chick embryos, we determined its spatio-temporal expression profile by whole-mount *in situ* hybridisation (WISH) and cryosectioning (Fig. 1A). At HH11-12, miR-128 was found in the neural tube, developing somites and in the notochord (Fig. 1i-i’). At HH16-17, miR-128 was detected in the myotome, with no expression apparent in the notochord and weak expression in the dorsal neural tube (Fig. 1ii-ii’’). At HH21-22, miR-128 was found in the branchial arches, around the eye and in fore- and hind limbs (Fig. 1iii-iii’’). Interestingly, miR-128 expression in the chick myotome is similar to the conserved microRNAs: miR-1, miR-133a/b and miR-206 (Ahmed et al., 2015; Sweetman et al., 2008).

**Figure 1:**
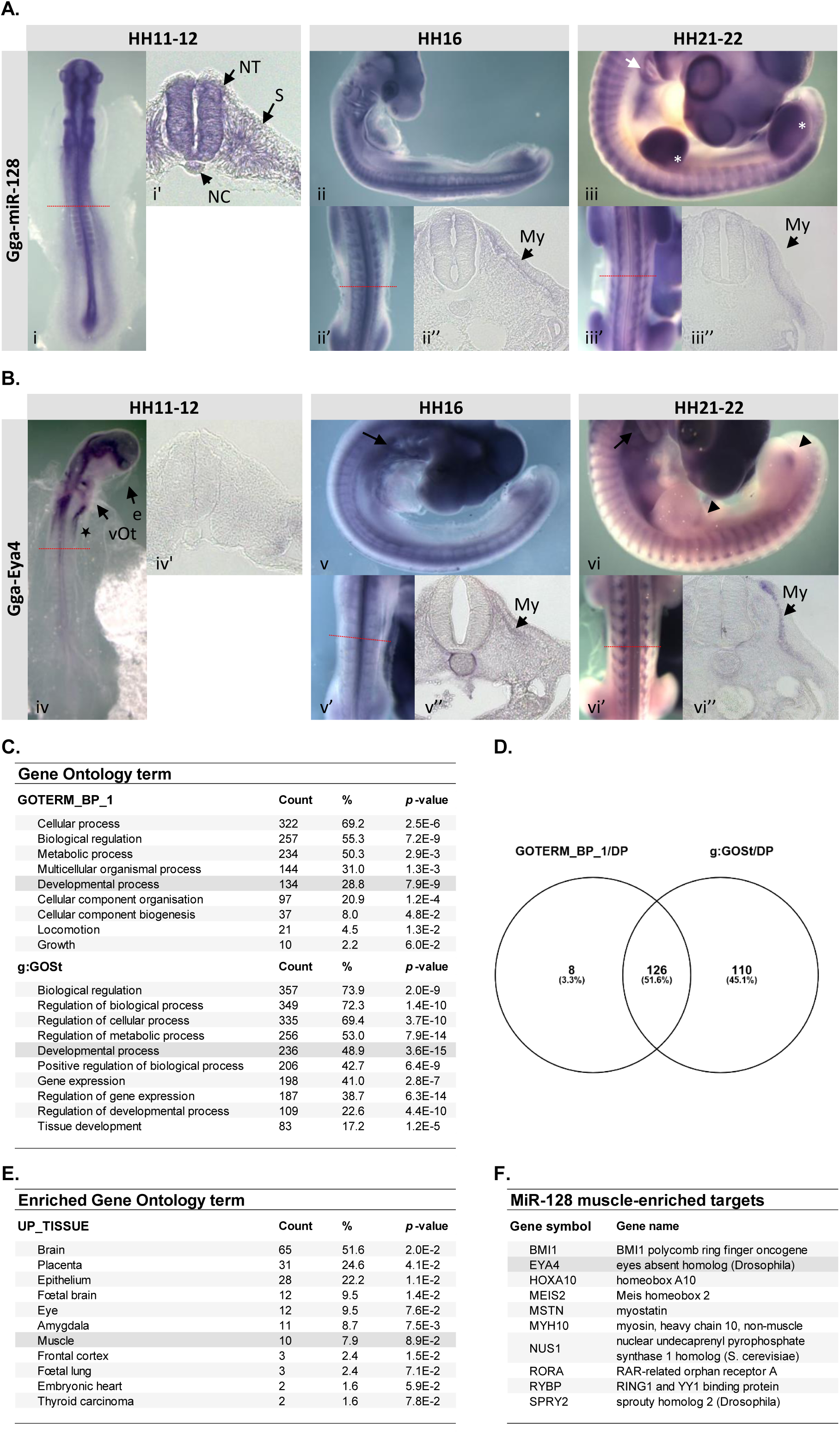
MiR-128 and *Eya4* are expressed in the myotome and many predicted miR-128 target genes are involved in developmental processes. Expression profiles of (A) miR-128 and (B) *Eya4* were determined by whole-mount in situ hybridisation at the stages indicated. The level of transverse sections at HH11-12 (i-i’)(iv-iv’), HH16 (ii-ii’’)(v-v”), and HH21-22 (iii-iii’’)(vi-vi”) are indicated by a red line. (A) At HH11-12, miR-128 is expressed in the neural tube (NT) and the developing somites (i’). At HH16 and HH21-22, miR-128 is in the branchial arches, in the myotome, and the developing limbs (ii-iii’’). At HH21-22, miR-128 is also expressed around the eye, and in the limbs (iii; white asterisk). (B) WMISH performed with antisense DIG-labelled RNA probe, and transverse sections at HH11-12, HH16, and HH21-22 At HH11-12 (iv-iv’), *Eya4* is expressed in the eye (e), the otic vesicle (vOt), and in a pool of non-identified migrating cells close to the heart region (iv; asterisk). At HH16 (v-v’’), *Eya4* is expressed in the eye, the branchial arches (v; arrow) and in the somites (s), in the myotome (v’’; My). At HH21-22 (vi-vi’’), *Eya4* is still expressed in the branchial arches (vi; arrow), and is strongly expressed in the myotome (vi’’). *Eya4* is also found dorsally in the posterior limb buds (vi; arrowhead). No expression was detected on embryos treated with the sense probe (negative control; data not shown). E: eye; My: myotome; NC: notochord; NT: neural tube; S: somite; vOt: otic vesicle. (C) Gene Ontology analysis of miR-128 muscle targets predicted by TargetScan. (D) Venn diagramme of the genes included in the category “Developmental process” using either GO term or g:GOSt. (E) For the 126 genes identified by both analyses the UP_TISSUE terms are shown. (F) The miR-128 muscle-enriched target genes (10) include *Eya4*.

To determine possible targets of miR-128 we used TargetScan (release 7.2; March 2018); this generated a list of 513 potential target genes. The molecular functions of miR-128 predicted targets was determined using GO term and g:GOSt analyses (Fig. 1C). The GO term annotation showed that 69.2% of miR-128-predicted targets were classified as ‘cellular process’. Other enriched GO terms included ‘biological regulation’ (55.3%) and ‘developmental process’ (28.8%). The g:GOSt analysis performed using g:Profiler showed similar results for broad categories. However, a larger number of targets were classified as playing a role in ‘developmental process’. The miR-128 targets listed in this category from GOTERM_BP_1 and g:GOSt were compared and 126 targets were common between the two different tools (Fig. 1D). Of these 126 targets more than 50% were found in the brain (65), which is notable as miR-128 was described as brain-enriched, about 10% were found in the eye (12), 8% in muscle (10) and less than 2% in the heart (2) (Fig. 1E).

One of the ten predicted targets for miR-128 in muscle was *EYA4* (Fig. 1F), a member of the PAX-SIX-EYA-DACH (PSED) network of transcriptional regulators, which act upstream of the MRFs in myogenesis. To examine a possible interaction between miR-128 and *EYA4* we characterised the expression profile of *EYA4* by whole mount in situ hybridisation (WISH) (Fig. 1B). At HH11-12, *EYA4* was mainly expressed in neural folds in the head region and in some cranial placodes, such as the optic and otic vesicles (Fig. 1Biv). *EYA4* was also expressed in a region proximal to the inflow region of the heart. At this stage no expression was detected in somites (Fig. 1Biv’). As the embryo developed, *EYA4* transcripts were detected in the branchial arches and somites. From HH16, *EYA4* was seen in the myotome with stronger expression at HH21-22 in the dorsomedial lip of the dermomyotome (Fig. 1Bv-vi’’). *EYA4* transcripts were also detected in dorsal root ganglia and in a posterior region of the developing limbs (Fig. 1Bvi). Thus, expression of miR-128 and *EYA4* was overlapping in the myotome and limb buds. We therefore examined whether miR-128 regulates *EYA4* expression post-transcriptionally and tested whether a direct interaction could be confirmed.

### Luciferase assays confirm negative regulation of *EYA4* by miR-128

To validate a potential interaction of miR-128 with the 3’UTR of *EYA4* (Fig. 2A), we generated luciferase reporters, both wild-type (WT) constructs and constructs where the miR-128 target site was mutated (mut). A potential miR-128 site was predicted in the 5’ part of the 3’UTR sequence by TargetScan and MiRanda algorithms; miR-27b has the same seed sequence as miR-128 and is predicted to target the same site (Fig. 2A, B). Additional candidate target sites in the *EYA4* 3’UTR included sites for miR-1a and miR-206, which have the same seed, as well as miR-133 (Fig. 2A, B; Fig. 3B). It has been found that miRNA sites located at the 5’ and 3’ extremities of a 3’UTR sequence are more likely to be functional (Long et al. 2007; Ekimler & Sahin 2014). Therefore, it is noteworthy that all these sites are located within the first 1,000 bp of the *EYA4* 3’UTR sequence, which is 6,000 bps in total. A 1kb fragment was cloned into a luciferase reporter and mutant constructs were generated for miR-27b/128, miR-1a/206 and 133a. The base pairing between the microRNAs and the putative target sites are shown and the mutated nucleotides are indicated (Fig. 2B).

**Figure 2:**
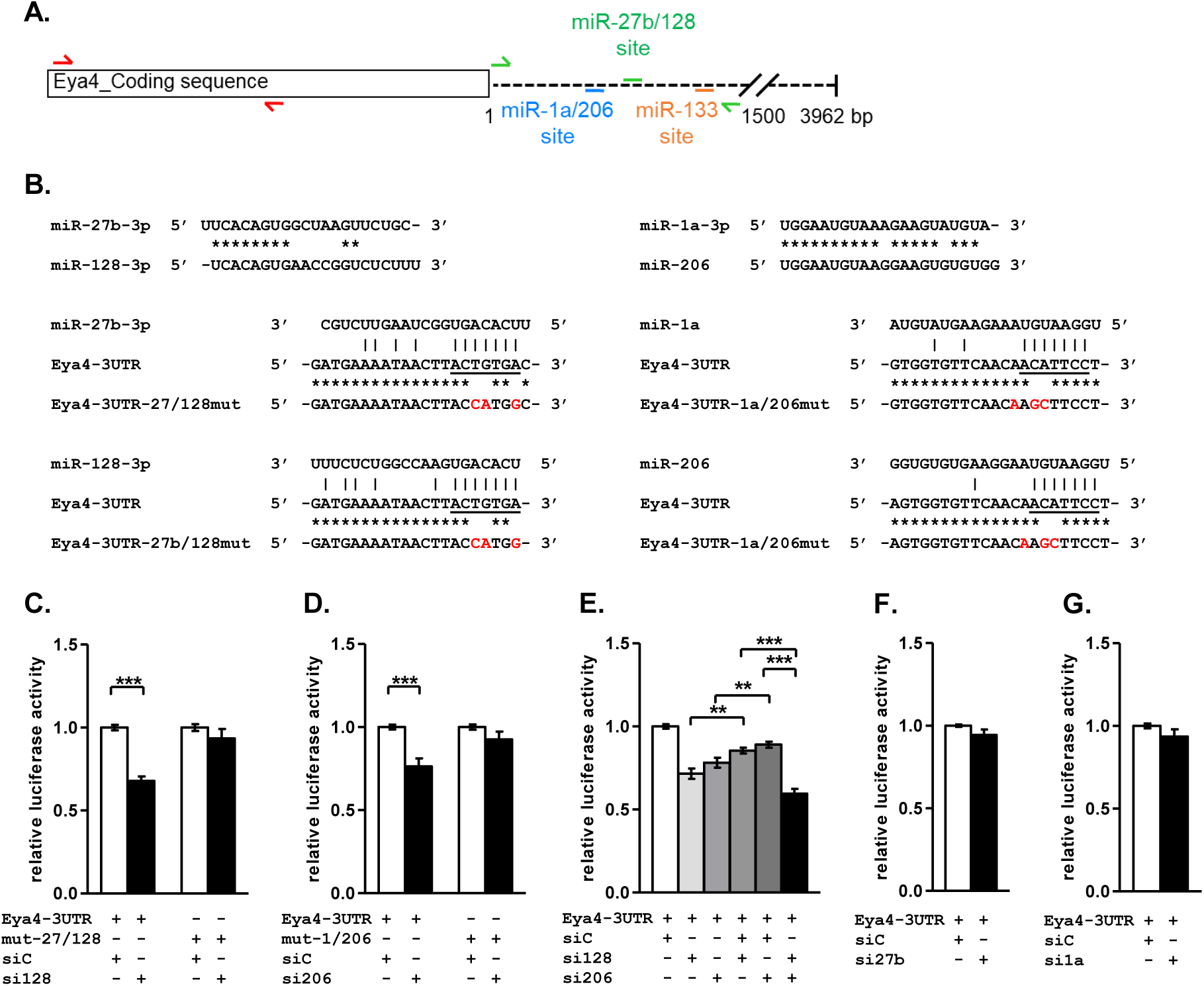
The *Eya4* 3’UTR contains functional target sites for miR-128 and miR-206. (A) Schematic of *Eya4* transcript with coding region (box) and 3’UTR (dotted line). Red and green arrows indicate the position of primer pairs used to clone the fragments of coding and 3’UTR sequences which were used for ISH and luciferase reporter assays, respectively. The positions of putative miR-1a/206, miR-27b/128 and miR-133 binding sites are indicated. (B) Alignment of miR-128 and miR-27b shows that the seed sequences are conserved but they have poor homology outside the seed. Alignment of miR-1a with miR-206 shows that they are very similar. Alignments of each of these microRNAs with their predicted target sites in the Eya4 3’UTR. Mutations introduced into the predicted target sites were designed to disrupt base pairing in the seed region (mutated nucleotides are indicated in red). Vertical lines indicate complementarity and asterisks indicate identity between sequences. (C-G) Relative Luciferase activity for Gga-Eya4 3’UTR reporter assays is shown, wild-type (WT) or mutants were co-transfected either with control siRNA (siC; white columns), or with mimics for miR-128 (C), miR-206 (D), miR-128 and miR-206 (E), miR-27b (F), or miR-1a (G) (black, grey shaded columns). Normalised luciferase activity was plotted relative to the siC condition. Experiments were repeated at least four times independently with triplicate samples in each. Error bars represent the standard error of the mean (SEM). Unpaired *t*-test: p<0.01: **, p<0.001: ***.

**Figure 3:**
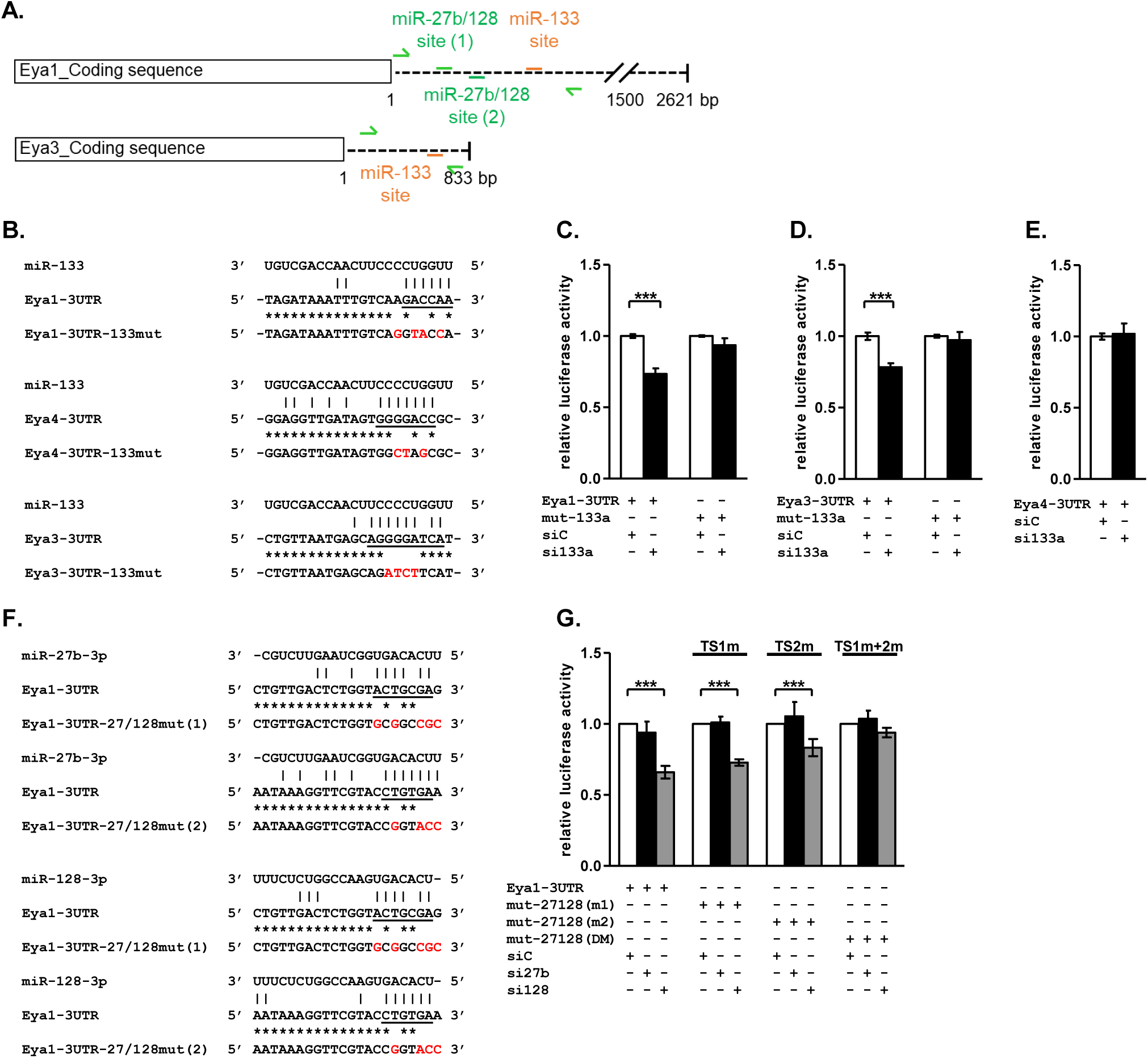
The *Eya1* and *Eya3* 3’UTRs contain functional target sites for miR-133a. (A) Schematic of *Eya1* and *Eya3* transcripts with coding region (box) and 3’UTR (dotted line). Green arrows indicate the position of primer pairs used to clone the fragments of 3’UTR sequences which were used for luciferase reporter assays. The positions of putative miR-27b/128 and miR-133 binding sites are indicated. (B) Alignments of *Eya1, Eya3* and *Eya4* 3’UTR predicted target sequences with miR-133a; mutated nucleotides disrupting seed-pairing in red. (C-E) Relative Luciferase activity for Gga-Eya1, or Gga-Eya3, or Gga-Eya4 3’UTR reporter assays is shown, wild-type (WT) or mutants were co-transfected either with control siRNA (siC; white columns), or with mimics for miR-133a (black columns). (F) Alignments of Gga-Eya1 3’UTR predicted miR-27b/miR-128 target sequences with miR-27b and miR-128 sequences. (G) Relative Luciferase activity for Gga-Eya1 3’UTR construct, wild-type (WT) and mutants (single and double mutants), co-transfected either with control siRNA (siC; white), or with mimics for miR-27b (black), or miR-128 (grey). Experiments were repeated 3 times independently with triplicate samples in each. Error bars represent the standard error of the mean (SEM). Unpaired *t*-test: p<0.001: ***.

In Luciferase assays co-transfection of miR-128 mimics with the *EYA4* 3’UTR lead to a decrease in relative luciferase activity compared to control (siC) (68% activity; t-test: p<0.001). Luciferase activity was restored to 93.5% of control by mutating the miR-128 binding site (Fig. 2C). Co-transfection of miR-206 mimics also regulated the *EYA4* 3’UTR. A decrease in luciferase activity was observed (76% activity; t-test: p<0.001), which was restored by mutating the miR-206 site (92.5% activity) (Fig. 2D). Interestingly, miR-27b did not affect luciferase expression suggesting it does not interact with the *EYA4* 3’UTR (Fig. 2F). Similarly, miR-1a did not interact with the *EYA4* 3’UTR (Fig. 2G) even though it shares the same seed sequence with miR-206. This indicates that miR-target gene interactions are not only based on complementarity between the seed and 3’UTR sequences and that additional nucleotides mediate specificity and need to be taken into account. Co-transfection of miR-133 mimic with the *EYA4* 3’UTR reporter had no effect (Fig. 3E).

Seed sequences are short (6-8 nts), and miRNAs often regulate several hundred targets. Conversely, multiple miRNAs can regulate the expression of a single gene by targeting different sites on the 3’UTR of its mRNA (Selbach et al. 2008; Bartel 2009). Because miR-128 and miR-206 led to a decrease in luciferase activity of the *EYA4* 3’UTR reporter we examined whether they cooperate. As before, luciferase activity was decreased in response to transfection of miR-128 mimic alone or miR-206 mimic alone (by 28.5% or 21.8%, Fig. 2E). This was very similar to the decreases observed in the previous experiments (32% or 24%, Fig. 2C, D). Smaller decreases in luciferase activity were observed with half the concentration of miR-128 or miR-206 mimic (mixed 1:1 with control mimic to give the same final concentration of oligo), 14.6% and 11% respectively. However, when miR-128 and miR-206 mimics were co-transfected expression of the *EYA4* 3’UTR luciferase reporter was reduced by 40.4% (Fig. 2E). Thus, their combined effect is greater than the sum of their individual effects, suggesting that miR-128 and miR-206 synergize (Ivanovska & Cleary 2008; Lu & Clark 2012).

### Members of the PSED network are regulated by myogenic microRNAs

Next, we investigated potential miRNA-mediated regulation of other members of the PSED network: *EYA1/3, SIX1/4* and *DACH1*. Using TargetScan and miRanda, several miRNAs were identified and predicted to target PSED members (Suppl. Table 1). Here, we focused on miR-128, miR-1/206 and miR-133 (Table 1; Figs. 3 and 4). Fragments of the 3’UTRs were cloned to generate luciferase reporter constructs, and mutations were introduced into the putative miRNA sites.

**Table.1.**
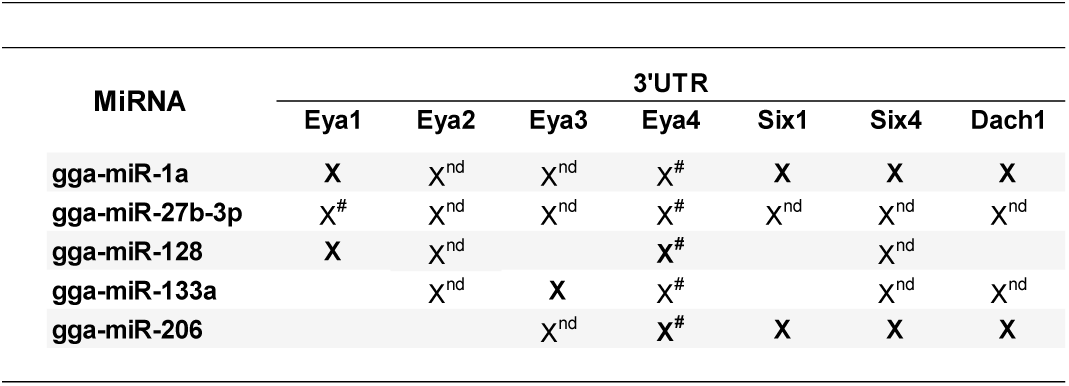
MiRanda analysis. Each miRNA was used to scan the 3’UTR sequences of chicken (Gallus gallus, gga) *Eya1* [ENSGALT000000025181.4], *Eya2* [ENSGALT00000007180.4], *Eya3* [ENSGALT00000001127.4], *Eya4* [ENSGALT00000022662.4], *Six1* [NM_001044685.1], predicted *Six4* [XM_003641442.2], and *Dach1* [ENSGALT00000027373.3]. #: MRE annotated in human sequence (TargetScan ‘human’), and conserved in chicken (TargetScan ‘chicken’). Predicted and validated targets are indicated in bold. nd: not studied in this work.

| MiRNA | 3'UTR |  |  |  |  |  |  |
| --- | --- | --- | --- | --- | --- | --- | --- |
|  | Eya1 | Eya2 | Eya3 | Eya4 | Six1 | Six4 | Dach1 |
| gga-miR-1a | X | X<sup>nd</sup> | X<sup>nd</sup> | X<sup>#</sup> | X | X | X |
| gga-miR-27b-3p | X<sup>#</sup> | X<sup>nd</sup> | X<sup>nd</sup> | X<sup>#</sup> | X<sup>nd</sup> | X<sup>nd</sup> | X<sup>nd</sup> |
| gga-miR-128 | X | X<sup>nd</sup> |  | X<sup>#</sup> |  | X<sup>nd</sup> |  |
| gga-miR-133a |  | X<sup>nd</sup> | X | X<sup>#</sup> |  | X<sup>nd</sup> | X<sup>nd</sup> |
| gga-miR-206 |  |  | X<sup>nd</sup> | X<sup>#</sup> | X | X | X |

**Figure 4:**
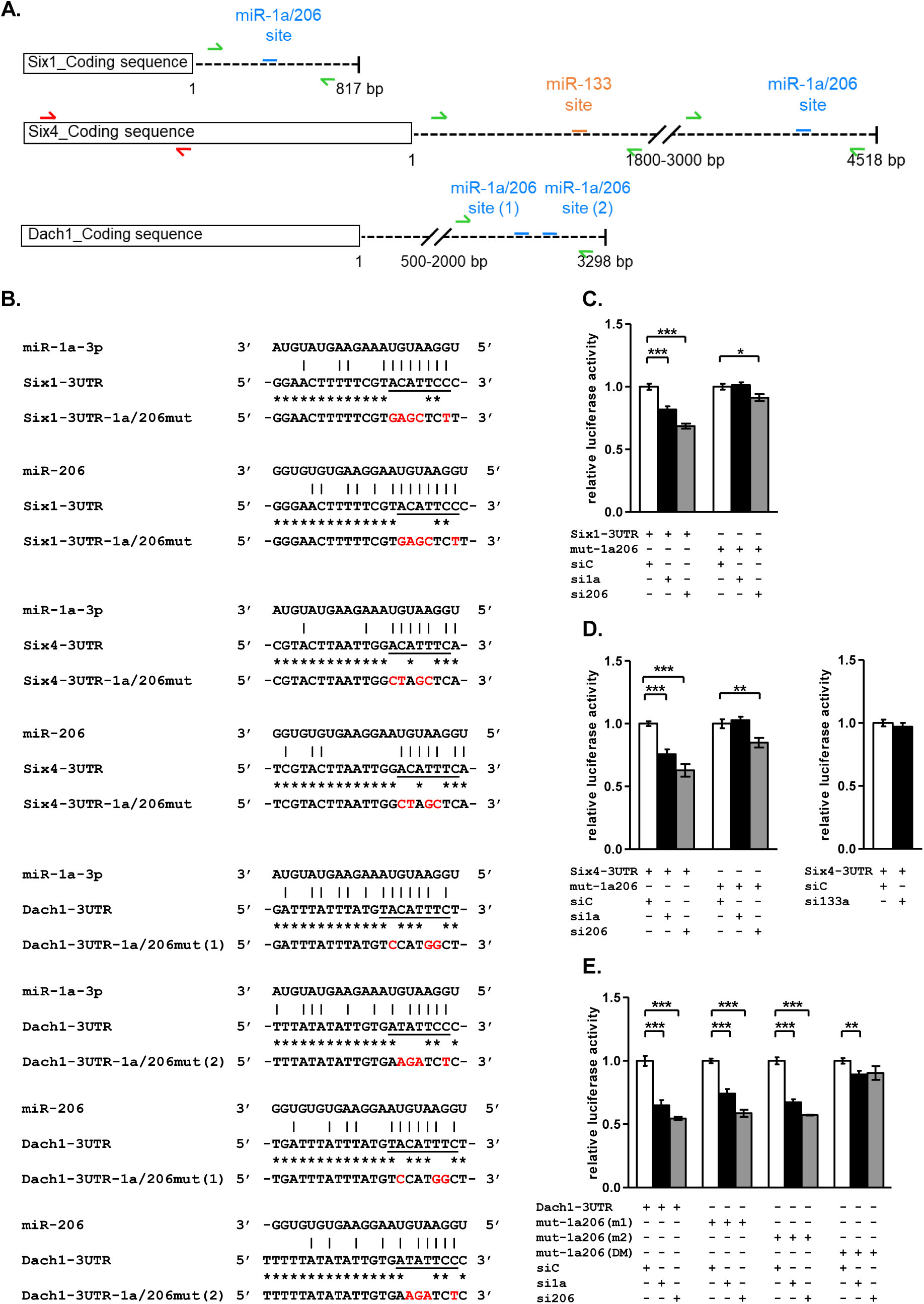
The PSED members *SIX1, SIX4* and *DACH1* are regulated by miR-1a/miR-206. (A) Schematic of *Six1, Six4* and *Dach1* transcripts with coding region (box) and 3’UTR (dotted line). Green arrows indicate the position of primer pairs used to clone 3’UTR fragments used for luciferase reporter assays. The positions of putative miR-1a/miR-206 and miR-133 binding sites are indicated. Red arrows indicate the position of primer pairs used to clone a *Six4* fragment used for ISH. (B) Alignments of *Six1, Six4* and *Dach1* 3’UTR predicted target sequences with miR-1a and miR-206; mutated nucleotides disrupting seed-pairing in red. (C-E) Relative Luciferase activity for Gga-Six1 (C), Gga-Six4 (D) and Gga-*Dach1* (D) 3’UTR constructs, wild-type (WT) and mutants, co-transfected either with control siRNA (siC; white), or with mimics for miR-1a (black) or miR-206 (grey). Experiments were repeated at least 3 times independently with triplicate samples in each. Error bars represent the standard error of the mean (SEM). Unpaired *t*-test: p>0.05.

We examined *EYA1* and *EYA3*, two additional EYA family members with putative target sites in their 3’UTRs (Fig. 3A, B). Luciferase reporter assays confirmed that miR-133 targets both *EYA1* and *EYA3*, but not *EYA4* (Fig. 3C-E). Specifically, luciferase activity of *EYA1* 3’UTR reporter was decreased by 26.6% (73.4% activity; t-test: p<0.001) in response to miR-133a mimic. This was restored by mutating the miR-133 site with relative luciferase activity going back to 94% (Fig. 3C). Luciferase reporter activity of *EYA3* 3’UTR decreased by 21.7% (78.3% activity; t-test: p<0.001) in response to miR-133a mimic. This was restored to 97.3% activity after mutation of miR-133 site (Fig. 3D). No effect on luciferase activity was observed using the *EYA4* 3’UTR reporter (Fig. 3E).

In addition, *EYA1* is regulated by miR-128 via two independent sites in the 3’UTR (Fig. 3F, G). With both putative target sites (TS) present in the *EYA1* 3’UTR, a decrease of 34% of the luciferase activity was observed in response to miR-128 mimic (66% activity). Mutation of each site individually (TS1, TS2) only partially restored luciferase activity levels, to 73% and 83% respectively. However, mutation of both sites restored luciferase activity to 94%. This shows that both TS1 and TS2 can work independently with TS2 being slightly more effective. Interestingly, the *EYA1* 3’UTR did not respond to miR-27b mimic.

Other members of the PSED network are *SIX1, SIX4* and *DACH1* and whilst there was no predicted target site for miR-128 in their 3’UTRs, TargetScan and miRanda predicted sites for miR-1a/206, miR-133 and miR-499 (Fig. 4A, B). Luciferase reporter assays showed a decrease in luciferase activity after transfection of mimics for either miR-1a or miR-206. For all confirmed target genes, *SIX1, SIX4* and *DACH1*, the effect of miR-206 was stronger compared to miR-1. Specifically, relative luciferase activity of the S*IX1* 3’UTR reporter decreased by 18.2% with miR-1 and by 31.4% with miR-206. For the *SIX4* 3’UTR reporter we observed a decrease of 24.3% with miR-1 and 37.2% with miR-206 (Fig. 4C, D). The *SIX4* 3’UTR reporter did not respond to miR-133 mimic (Fig. 4D), or to miR-499 mimic (not shown).

The *DACH1* 3’UTR reporter contained two predicted target sites for miR-1a and miR-206. Transfection of mimics for these microRNAs led to a decrease of 35% and 45.4% in luciferase activity, respectively. Introducing point mutations into the miR-1a/206 sites separately only led to a minor rescue of luciferase activity to 74% and 59% activity respectively (t-test: p<0.001) (Fig. 4E), suggesting that the two sites can work independently. Mutation of both sites restored luciferase activity to approximately 90%.

### Myogenesis is impaired after miR-128 knock-down and the PSED network - including EYA4 - is deregulated

Given the restricted expression of miR-128 in the somite myotome, we asked whether miR-128 is required for myogenesis *in vivo*. In addition, we determined whether miR-128 loss-of-function had an effect on expression of the PSED network and in particular on *EYA4* (Fig. 5). The most posterior six somites of HH14-15 chicken embryos were injected with antagomir-128 (AM-128), or with a scrambled antagomir (AM-scr). The resulting phenotypes were assessed after 24 hours using whole mount *in situ* hybridisation and cryosections (Fig. 5A-C) or RT-qPCR (Fig. 5D). Non-injected contralateral somites were used as additional controls.

**Figure 5:**
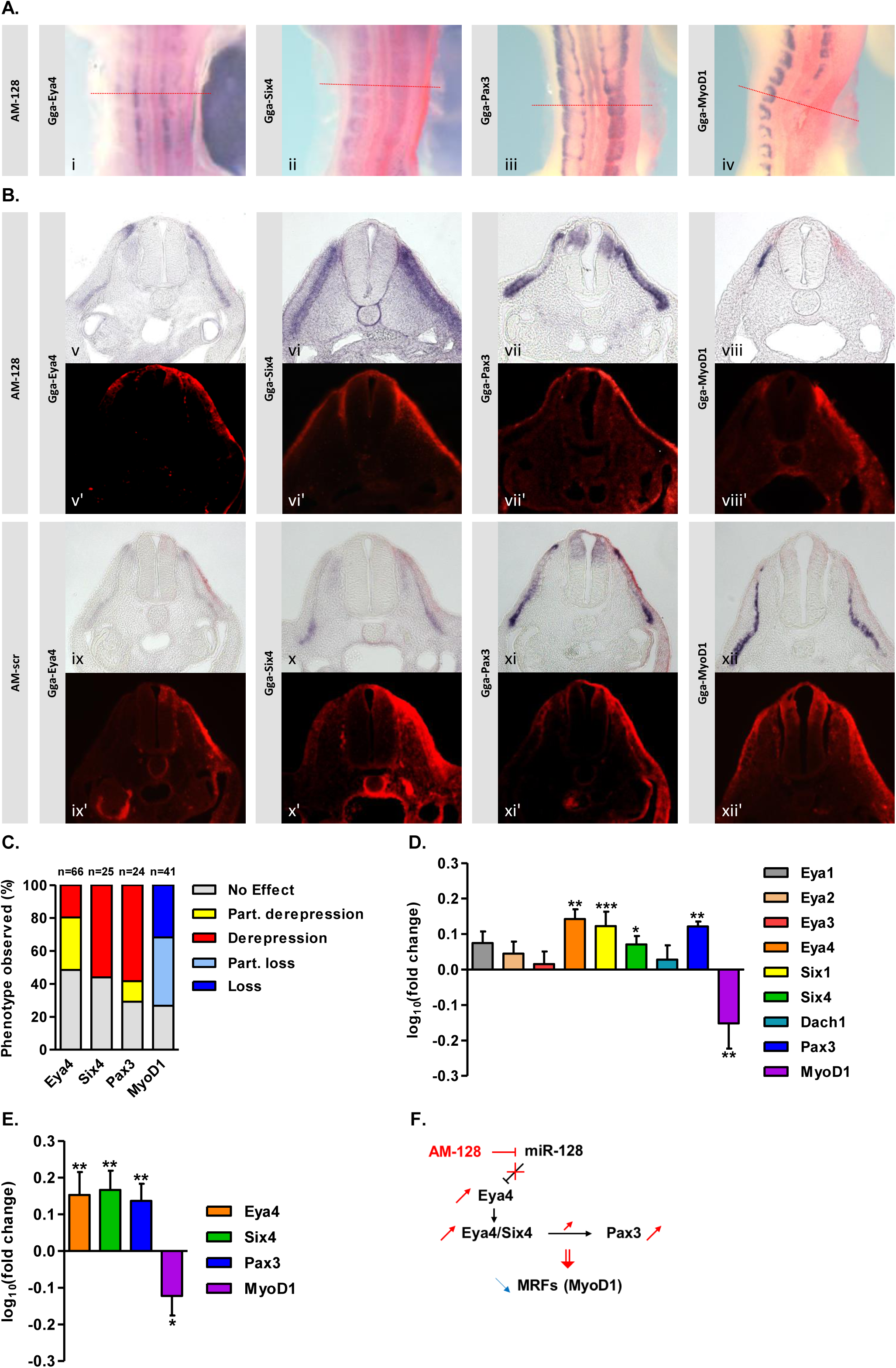
MiR-128 regulates myogenesis through its interaction with *Eya4*. AntagomiR-128 (1 mM) was injected into the 6 most posterior somites, on one side of HH13-14 embryos. After 24h incubation, HH19-20 embryos were processed for ISH to detect the transcripts indicated. To identify injected somites the FITC-coupled antagomir-128 or antagomir-scr was detected using Fast Red. (A) Expression patterns in whole mount embryos, dorsal view. (B) Transverse sections (red dotted lines) at the interlimb level showing *Eya4, Six4, Pax3* and *MyoD1* expression as indicated; Alexa-Fluor-568 reveals antagomiR location. The contralateral non-injected side (left side), was used as control. (C) Quantification of phenotypes observed. (D) PSED members and MRFs (MyoD1 and Myf5) transcript levels in somites injected with AM-128. RT-qPCR results expressed in log_10_(fold change). For each gene and each experiment, the injected somite data were normalised to two housekeeping genes (GAPDH+β-actin) and compared to the contralateral non-injected somite data of the same embryo. Number of independent experiments for each tested gene after AM-128 injection: *Dach1* (n=6); *Eya4, Six4, Pax3, MyoD1* (n=7); *Eya3, Six1* (n=8); *Eya1, Eya2* (n=10). (E) For targeted mis-expression Eya4 expression construct was injected and electroporated into the 6 most posterior somites, on one side of HH13-14 embryos. After 24h incubation, HH19-20 embryos were collected, injected somites were dissected for RNA extraction. RT-qPCR showed the deregulation of PSED members (*Eya4, Six4, Pax3*) and *MyoD1* transcripts as (log_10_(fold change)) in electroporated somites compared to controls. At least seven independent experiments were performed for each gene. Mann-Whitney *U*-test: p<0.05: *, p<0.01: **, p<0.001: ***. (F) Summary: Inhibiting the negative regulation of Eya4 by miR-128 led to de-repression of Eya4, which together with Six4 activates expression of Pax3. This prevents entry of myogenic progenitors into the differentiation programme, indicated by loss of MyoD1 expression.

Antagomir-mediated KD of miR-128 led to decreased *MYOD1* expression in developing somites. This was clearly visible in whole mount (Fig. 5A). Cryosections confirmed the loss of *MYOD1* transcripts detected in the myotome (Fig. 5B). Grouping the ISH results showed that the majority of embryos (73.2%) had either complete (n=13/41, 32%) or partial loss of *MYOD1* (n=17/41, 41%). The qualitative ISH data was confirmed by RT-qPCR of dissected somites, which were pooled from seven independent injection experiments. This also showed a reduction of *MYOD1* expression (Fig. 5D).

Interestingly, miR-128 KD led to a concomitant increase in expression of the pro-myogenic transcription factor and PSED member, *PAX3*, which showed an increase in the dermomyotome, especially in its central part where its expression is usually weak (Fig. 5B). The majority of embryos (70.8%) showed an increase of *Pax3*, 14/24 embryos were similar to the embryo shown (Fig. 5Aiii, 5Bvii), 3/24 showed a partial increase and 7 embryos showed no change. A relative increase in *Pax3* transcript levels (30%) after AM-128 injection into somites was confirmed by RT-qPCR (1.3-fold change; t-test: p<0.01) (Fig. 5A-D). This indicates that myogenic cells in the dermomyotome remain in a progenitor stage of development and did not activate the differentiation programme (Williams & Ordahl 1994; Goulding et al. 1994; Gros et al. 2004).

In whole-mount embryos, expression of *EYA4*, a validated direct target of miR-128, did not seem to be different between AM-128 injected and non-injected sides (Fig. 5Ai). However, transverse cryosections showed that *EYA4* expression was increased in the central part of the myotome after miR-128 KD (Fig. 5Bv). This phenotype was observed in 51.5% of the embryos: 13/66 were similar to the embryo shown, 21/66 embryos showed partial de-repression (31.8%) and 48.5% of the embryos (n=32) showed no change in expression on the injected side compared to the non-injected side. RT-qPCR confirmed the increase in *EYA4* expression, which was 1.4-fold higher in pooled somites injected with AM-128 compared to non-injected pooled somites from the opposite side (t-test: p<0.01) (Fig. 5D). Thus, ISH and RT-qPCR showed that KD of miR-128 resulted in de-repression of *EYA4* expression in the myotome. This is consistent with the *in vitro* luciferase reporter experiments (Fig. 2B) and identifies *EYA4* as a direct miR-128 target *in vivo*.

To examine potential effects of miR-128 KD on other members of the PSED network we used RT-qPCR to assess the expression of *EYA1, EYA2, EYA3, SIX1, SIX4* and *DACH1* (Fig. 5D). In addition, we performed WISH for SIX4 (Fig. 5A, B). The expression of *EYA1* was 1.2-fold higher than the control, but *EYA2* and *EYA3* were not affected after AM-128 injection (t-test: p<0.05) (Fig. 5D). The increase of *EYA1* is consistent with luciferase reporter assays, which identified *EYA1* as a direct target for miR-128 *in vitro* (Fig. 3F, G). Although *SIX1* and *SIX4* were not predicted as direct targets of miR-128, a 1.3-fold increase in their expression levels was observed after antagomiR-128 injection, most likely due to indirect effects. ISH performed on AM-128 injected embryos also showed an increase in *SIX4* expression in injected somites, in the dorsomedial lip of the dermomyotome and the myotome (Fig. 5Aii, 5Bvi). This was observed in 14/25 embryos (56%) (Fig. 5C). These results are consistent with a potential indirect effect and cross-regulation between SIX and EYA co-factors, which have been shown to form a strong complex that activates SIX target genes (Grifone et al., 2005; Heanue et al., 1999; Ohto et al., 1999).

Next we asked whether targeted mis-expression of EYA4 would mimic the miR-128 KD phenotype. A pCAβ-*Gga-Eya4*-full-length expression construct was injected and electroporated into posterior somites of HH14-15 chicken embryos. After 24h, embryos were harvested and successfully electroporated somites (GFP^+ve^) and non-injected contralateral somites were dissected for RT-qPCR. This confirmed a 1.53-fold increase in *EYA4* expression and a concomitant 1.54- and 1.44-fold increase of *SIX4* and *PAX3* expression (n=7-9; t-test: p<0.01) (Fig. 5E). The increased expression of the PSED members correlated with 0.8-fold decrease of *MYOD1* expression (n=6; t-test: p<0.05). Therefore, elevated expression of EYA4, either due to miR-128 KD or due to electroporation led to deregulation of the PSED network and inhibition of myogenesis *in vivo*, suggesting that fine-tuning the expression of EYA4 in the myotome by miR-128 contributes to control the entry into the myogenic program (Fig. 5F).

## Discussion

The MRFS control entry into the myogenic program, which leads to the formation of skeletal muscle. Upstream of this obligatory step signalling molecules and other transcription factors, including the PSED network, direct cells toward myogenesis. Therefore, members of this regulatory network, PAX, SIX, EYA and DACH are referred to as pre-myogenic factors. In vertebrates, SIX1/4, EYA1/2/4, and DACH1/2, have overlapping expression patterns in the myogenic precursor cells in the somites, in the dermomyotome and the myotome. At limb level, together with PAX3, they play a crucial role in ensuring that migrating myogenic precursor cells remain committed to their fate until they reach their final destination (Christ and Ordahl, 1995).

Experiments in mice and chicken provided insight into the roles of PSED members upstream of the activation of the myogenic program. Ectopic expression of PAX3, SIX or EYA in chicken embryos led to activation of PAX3 and MRFs (Heanue et al., 1999; Maroto et al., 1997). While in *PAX7*^*-/-*^ mutant mice skeletal muscle forms normally (Mansouri et al., 1996), *PAX3*^*-/-*^ mutants have abnormal myotome formation, trunk muscle defects and absence of limb muscle (Bober et al., 1994; Goulding et al., 1994). Moreover, *PAX3*^*-/-*^*/PAX7*^*-/-*^ double-mutant mice have major defects in myogenesis (Relaix et al., 2005). Similarly, no developmental defects were observed in *SIX4*^*-/-*^ and *EYA2*^*-/-*^ mice (Grifone et al., 2007; Ozaki et al., 2001), but *SIX1*^*-/-*^ and *EYA1*^*-/-*^ mutant mice have important muscle deficiencies (Grifone et al., 2007; Laclef et al., 2003; Ozaki et al., 2004), while *SIX1*^*-/-*^*/SIX4*^*-/-*^ and *EYA1*^*-/-*^*/EYA2*^*-/-*^ double-mutant mice lack all muscles derived from the hypaxial dermomyotome. *SIX1*^*-/-*^*/SIX4*^*-/-*^*/MYF5*^*-/-*^ triple-mutant mice display a similar phenotype to what was observed in *PAX3*^*-/-*^*/MYF5*^*-/-*^ mutants, with no expression of MYOD and no skeletal muscle formed (Giordani et al., 2007; Relaix et al., 2013; Tajbakhsh et al., 1997), suggesting that SIX and EYA are upstream of PAX.

Transcription regulation of target genes by SIX proteins requires cooperative interaction with EYA proteins (Ohto et al., 1999), moreover SIX1/4 binding to EYA1/2 in the cytoplasm preceeds translocation into the nucleus. SIX, often associated with DACH, has been described as a repressor or weak activator, however, when interacting with EYA, the complex formed becomes a strong activator, which is then able to activate SIX target genes, such as *PAX3* and *MYOD1* and therefore influence myogenic differentiation (Grifone et al., 2005; Heanue et al., 1999). In addition, it has also been shown that SIX/EYA complex can directly up-regulate *MYOD* and *MYOG* expression by targeting enhancer elements on their respective promoters (Giordani et al., 2007; Spitz et al., 1998; Tapscott, 2005). These results are consistent with the severe decrease of *MYF5* and *MYOD1* expression, in the myotome, observed in the *SIX1*^*-/-*^ */SIX4*^*-/-*^ double-mutant mice (Buckingham and Rigby, 2014; Grifone et al., 2005).

Here we demonstrate the co-regulation of PSED members by multiple microRNAs that are known to be enriched in skeletal muscle (Mok et al., 2017). In particular, antagomir-mediated KD of miR-128 inhibits myogenesis, MYOD1 expression is lost on the injected side and expression of the premyogenic genes, PAX3, SIX4 and EYA4, is increased (Fig. 5A, B). We propose that the inhibition of myogenesis results from de-repression of *EYA4*, which we show is directly targeted by miR-128 via a site in the 3’UTR (Fig. 2C). The data suggest that increased level of EYA4 is keeping cells in a pre-myogenic state, potentially through the formation of transcriptional complexes with SIX proteins (SIX1/4), which then lead to increased expression of PAX3. This is illustrated in the model shown (Fig. 5F). Deregulation of EYA4 *in vivo*, in developing somites, also affects expression of other members of the PSED network as shown by RT-qPCR (Fig. 5D). The observed increase in *PAX3* expression is consistent with the fact that PAX3 is a known target of SIX (Grifone et al., 2005) Interestingly, a miR-128 site was predicted by TargetScan in the human PAX3 3’UTR. Thus, it is possible that *PAX3* is a direct target of miR-128 in human, however, a canonical target site (Bartel, 2009) was not identified in the chicken *PAX3* 3’UTR by any of the algorithms we used in this work.

Overexpression of EYA4 could phenocopy the miR-128 KD (Fig. 5E). This is consistent with the idea that *EYA4* is an important miR-128 target. Furthermore, both *EYA4* and miR128 are co-expressed in the myotome (Fig. 1). Luciferase assays also confirmed that miR-206 can negatively regulate the 3’UTR of *EYA4* and that miR-128 and miR-206 act cooperatively (Fig. 2). However, the highly related microRNAs miR-27b and miR-1 did not affect expression of the *EYA4* 3’UTR luciferase reporter (Fig. 2 F, G). These microRNAs share seed sequences with miR-128 and miR-206 respectively. miR-206 is also expressed in the myotome and we showed previously that it regulates *PAX3* thereby regulating the myogenic progenitor to committed myoblast transition (Goljanek-Whysall et al., 2011). This raises the possibility that miR-128 and miR-206 also cooperate *in vivo* targeting multiple members of the PSED network, including *EYA4* and *PAX3*, via direct and indirect mechanisms.

To determine if other PSED members might be regulated by microRNAs known to be involved in myogenesis, we identified potential target sites for miR-128, miR-27b, miR-1, miR-206 and miR-133 in the 3’UTRs of *EYA1, EYA3, SIX1, SIX4* and *DACH1*. Luciferease reporters showed that *EYA1* has two target sites that can be recognized by miR-128 but not by the related miR-27b (Fig. 3A, G). *EYA1* and *EYA3* each have one target site for miR-133 (Fig. 3A, C, D) and *SIX1* and *SIX4* each have one target site that is recognized by both miR-1 and miR-206. Furthermore, *DACH1* has two sites recognized by both miR-1 and miR-206 and these seem to work independently of each other.

Overall our data suggest that fine-tuning levels of the PSED transcriptional regulators in developing somites by multiple microRNAs, miR-128, miR-1, miR-206 and miR-133, is important for the myogenic differentiation program. In addition, we identify miR-128 as a novel microRNA required for myogenesis and EYA4 as an important direct target.

## Material and Methods

### Culture and staging of embryos

Fertilised White Leghorn chicken eggs (Henry Stewart & Co Ltd, UK) were incubated at 38°C until they reached the desired stage of development according to (Hamburger and Hamilton, 1951).

### Probes, *in situ* hybridisation, sections and photography

Fragments of chicken *Eya4* and *Six4* coding sequences were PCR amplified from embryonic cDNA using primers for *Gga-Eya4* [NM_001305177.1] and *Gga-Six4* [XM_003641442.2], cloned into pGEM-T Easy vector (Promega) and validated by sequencing. Antisense Digoxigenin-labelled RNA probes were generated for whole-mount *in situ* hybridisations as described (Mok et al., 2018). Double-DIG-labelled LNA oligonucleotide-containing probes for miR-128 (Exiqon) were used as described (Ahmed et al., 2015). After colour reaction, embryos were de-stained in 5X TBST detergent mix. Photography on a Zeiss SV11 stereo-microscope used QCapture software. For cryosectioning, PFA-fixed embryos were embedded in O.C.T., 20 µm sections were collected on SuperFrost-Plus slides, mounted with Hydromount and photographed on a Zeiss AxioPlan microscope using AxioVision software.

### DNA constructs, transfections and luciferase assay

Chicken *EYA1, EYA3, EYA4, SIX1, SIX4* and *DACH1* 3’UTR fragments containing predicted binding sites of miR-1a, miR-206 and mir-133a, miR-27b and miR-128 were PCR amplified and sub-cloned downstream of the luciferase gene as before (Goljanek-Whysall et al., 2014). Mutant constructs were generated using FastCloning; miRNA target sites predicted by TargetScan and MiRanda algorithms (Agarwal et al., 2015; Betel et al., 2008) were replaced with restriction enzyme sites introducing point mutations. All constructs were validated by sequencing.

Chick dermal fibroblast (DF1) were seeded into 96-well plates at 7×10^4^ cells/cm^2^ and transfected in triplicate with Renilla and firefly luciferase reporter plasmids (25 ng, 100 ng) with miRNA mimics, identical to endogenous mature miRNAs, or si-control (siC) (50 nM, Sigma) using Lipofectamine 2000 (Invitrogen). After 24 hours luciferase activities were assayed in cell lysates using the Dual-Luciferase Reporter Assay System (Promega) and a multi-label counter (Promega GloMax). Relative luciferase activities were obtained by calculating the ratios of Firefly to Renilla luciferase activity, which was normalised to siC-treated samples.

### Cloning of chicken *EYA4*, injection and electroporation into somites

The full-length coding region of chicken *EYA4* was PCR amplified from HH19-20 somite cDNA using SuperScript III Reverse Transcriptase kit (Invitrogen). Primer design used predicted *EYA4* sequences for chicken (ENSGALG00000031656) available from Ensembl Genome Browser (www.ensembl.org). PCR products were validated by sequencing and subcloned into the pCAβ expression vector.

Eggs were windowed and black ink was injected underneath the blastoderm to visualise the embryos. AntagomiRs AM-128 and scrambled antagomiR (AM-scr) (Dharmacon) were designed as described (Goljanek-Whysall et al., 2011) and injected into the posterior six somites of HH14-15 embryos, final concentration 1mM. After 24h embryos were harvested and processed for *in situ* hybridisation, or injected somites were dissected and processed for RNA extraction. Corresponding somites from the non-injected side were collected and used as control.

Expression construct (pCAβ-*Gga-Eya4*, 2 mg/mL) was injected into the posterior six somites of HH14-15 embryos and electroporated using five 20 ms pulses of 50 V (Sweetman et al., 2008). Plasmids produce GFP for tracing. Embryos were harvested after 24 h and those showing GFP fluorescence in targeted somites were processed.

### RNA extraction and RT-qPCR

TRIzol reagent (Ambion) was used to isolate RNA from somites according to manufacturer’s instructions; RNA was DNase treated (Roche) and extracted using acid phenol-chloroform (Ambion). cDNA was synthesised using random hexamer primers (Invitrogen) and SuperScript II Reverse Transcriptase kit (Invitrogen). RT-qPCRs were performed in 96-well plates on ABI Prism 7500 (Applied Biosystems) using SYBR Green according to manufacturer’s instructions. Primers (Sigma) were designed with PrimeTime (https://eu.idtdna.com/scitools/Applications/RealTimePCR/). Relative quantifications were calculated using the Relative Standard Curve method and normalised to the averaged relative quantification of *β-actin* and *GAPDH* housekeeping genes. Results from injected somites were compared to their contralateral non-injected somites, expressed in log_10_(fold change) and plotted on a linear scale where the x-axis corresponds to the non-injected condition set at 0 (log_10_(1)=0).

### Computational methods

MiRNA sequences were collected from XenmiR, GEISHA and miRBase databases (Ahmed et al., 2015) (Kozomara and Griffiths-Jones, 2014). Potential miRNA targets were identified using TargetScan (Agarwal et al., 2015; Lewis et al., 2005). Identification of potential miRNAs targeting mRNAs of interest was done using miRanda (Betel et al., 2008; John et al., 2004). GO term analysis was assessed using DAVID bioinformatics resources and g:Profiler.

### Statistical analysis

Luciferase assays and RT-qPCR data were analysed using Student’s two-tailed unpaired *t*-test and Mann Whitney *U*-test to assess the differences in one variable between non-treated and treated samples. The data are presented as the means ± S.E.M. unless indicated and are representative of at least three independent experiments. In all statistical analysis, p<0.05 was considered significant.

## Acknowledgements

We thank Timothy Grocott for help with cloning and luciferase assays and for discussions, Gi Fay Mok for help with microinjection and electroporation, Tracey Swingler for help with qPCR, Simon Moxon for help with bioinformatics. CV was funded by the BBSRC doctoral training programme on the Norwich Research Park. Research was supported by BBSRC project grants to AM (BB/K003437, BB/N007034).

